# Focus on single gene effects limits discovery and interpretation of complex trait-associated variants

**DOI:** 10.1101/2025.06.06.658175

**Authors:** Kathryn Lawrence, Tami Gjorgjieva, Stephen B. Montgomery

## Abstract

Standard QTL mapping approaches consider variant effects on a single gene at a time, despite abundant evidence for allelic pleiotropy, where a single variant can affect multiple genes simultaneously. While allelic pleiotropy describes variant effects on both local and distal genes or a mixture of molecular effects on a single gene, here we specifically investigate allelic expression “proxitropy”: where a single variant influences the expression of multiple, neighboring genes. We introduce a multi-gene eQTL mapping framework—*cis*-principal component expression QTL (*cis*-pc eQTL or pcQTL)—to identify variants associated with shared axes of expression variation across a cluster of neighboring genes. We perform pcQTL mapping in 13 GTEx human tissues and discover novel loci undetected by single-gene approaches. In total, we identify an average of 1396 pcQTLs/tissue, 27% of which were not discovered by single-gene methods. These novel pcQTL colocalized with an additional 142 GWAS trait-associated variants and increased the number of colocalizations by 34% over single-gene QTL mapping. These findings highlight that moving beyond single-gene-at-a-time approaches toward multi-gene methods can offer a more comprehensive view of gene regulation and complex trait-associated variation.

## Introduction

Co-expression of nearby genes is a widespread phenomenon. Empirically, across human tissues in GTEx, 13% to 53% of genes have expression correlated with their neighbor ^1^. This observed correlation can result from several biological mechanisms—for instance: transcription factors co-regulating multiple genes in *trans*, shared proximal regulatory elements like promoters and enhancers co-regulating multiple genes in *cis*, genes sharing a local chromatin state or epigenetic marks, etc.—or technical artifacts ^2–5^. Previous work has also revealed abundant allelic pleiotropy, where one variant associates with the molecular phenotypes of multiple nearby genes ^6^. Combined, these observations indicate that a proportion of non-coding genetic effects do not act on only one causal gene.

However, current approaches to understand the effects of genetic variation still focus on one-variant-one-gene mapping approaches, despite observations of co-expression and of sharing of *cis*-regulatory mechanisms among neighboring genes ^7^. For instance, the widely-utilized expression quantitative trait loci (eQTL) framework considers only one gene at a time where the expression of a single gene is regressed on a genetic variant, resulting in an estimate of that genetic variant’s linear effect on gene expression ^8^. We hypothesize that QTL methods jointly considering neighboring genes will improve our ability to detect and interpret the impact of common genetic variation on gene expression (Fig.1A).

**Figure 1.**
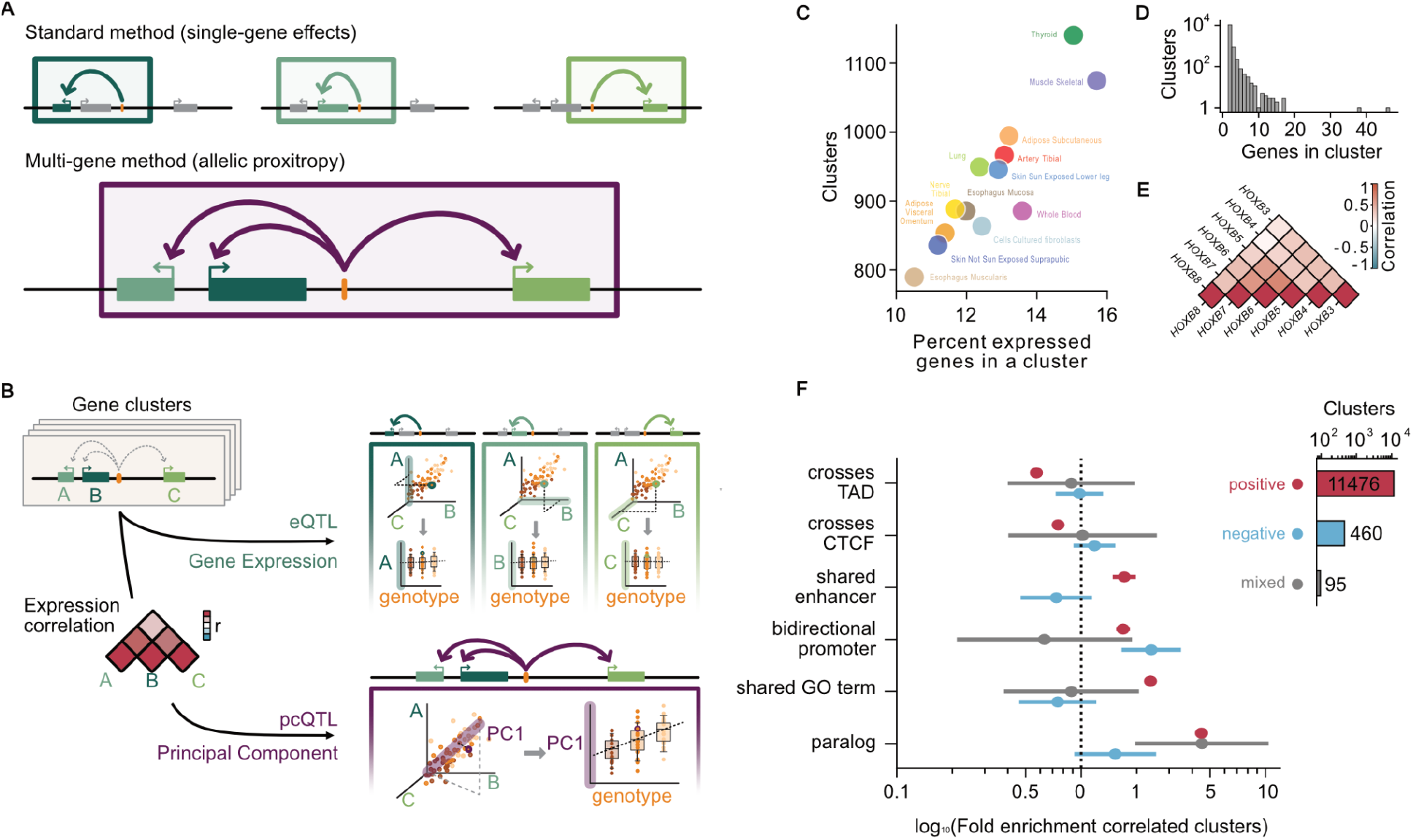
Multi-gene QTL mapping from correlated gene clusters. **A** Illustration showing standard single-gene methods detecting single-gene effects vs a multi-gene method detecting allelic proxitropy. **B** Illustration of the pcQTL pipeline. For each cluster of genes with correlated expression, eQTLs are equivalent to testing for correlation with genotype along the A, B, or C axis, while pcQTLs test for correlation with genotype along a constructed principal component axis. **C** The number of clusters in each tissue vs the percent of all expressed genes in the tissue that are in a cluster. **D** Distribution of the number of genes per cluster. **E** Example heatmap of a cluster of six *HOXB* genes with correlated expression in esophagus muscularis, colored by Spearman correlation coefficient. **F** Enrichment of correlated clusters split for clusters with all positive correlation, all negative correlation, or a mix of positive and negative correlation. Error bars are 95% CIs on odds ratios for logistic regression.

To overcome the limitations of single-gene analyses, we introduce a multi-gene QTL mapping approach—*cis*-principal component QTL (*cis-*pc eQTLs or pcQTLs for brevity)—to jointly analyze clusters of neighboring, co-expressed genes across 13 human tissues from GTEx (Fig.1B). Using this approach, we discover novel genetic effects missed by single-gene analyses. We further demonstrate improvements to colocalization with GWAS hits, uncovering 34% additional trait-associated genetic variants missed by the traditional single-gene eQTL approach. Our results demonstrate that jointly analyzing neighboring co-expressed genes leverages shared regulatory architecture and allelic proxitropy, improving our ability to detect and interpret genetic effects on gene expression and complex human traits.

## Results

### Transcriptome-wide identification of clusters of co-expressed neighboring genes

We started by identifying clusters of co-expressed neighboring genes (“gene clusters”) across human tissues using GTEx data. We focused on 13 tissues with large sample sizes (N>400) as these are best powered for eQTL analyses. As gene expression data often has global structured variance due to technical and known biological covariates, latent factor correction using tools like surrogate variable analysis ^9^, global principle component analysis (PCA) or probabilistic estimation of expression residuals (PEER) ^10^ are often employed to improve power to discover *cis*-regulatory effects ^8^. We found residualization of latent factors (60 PEER factors) also improved detection of co-expression of neighboring genes in GTEx data. While the un-residualized data showed frequent long-range correlations (median gene-gene distance 40 Mb), after residualization significant correlations are at the smaller scale of shared *cis*-regulation (median gene-gene distance 290 kb) (Supplementary Fig.1). This suggests latent factor correction enhances the accuracy of co-expression detection among neighboring genes by eliminating spurious long-range interactions, thereby highlighting significant correlations driven by shared *cis*-regulatory mechanisms at a localized scale.

We performed a transcriptome-wide analysis to identify clusters of nearby correlated genes; we did this for each tissue independently (Methods) This resulted in 787-1138 clusters across tissues—a total of 12,022 clusters—with between 10.5-15.7% of expressed genes in each tissue belonging to a cluster (Fig.1C; Supplementary Fig.2). The majority of clusters (89.2%) are pairs, 7.2% have 3 genes, and 3.6% have 4 or more genes (Fig.1D). We observed that it is more frequent for genes to be positively rather than negatively correlated, though 4.6% of all clusters across tissues (n=555) contain at least one pair of genes with a significant negative correlation (Fig.1F). Large clusters often represent known functionally-related groups of genes, such as the Hox cluster of 6 genes on chromosome 17 or the Keratin gene cluster (with 38 genes) on chromosome 17 (Fig.1E; Supplementary Fig.2). We further found that positively correlated gene clusters are enriched for similar biological functions and regulatory architecture. They are more likely to include paralogs, belong to the same gene ontology (GO) term, and have shared enhancers, but they are less likely to cross a topologically associated domain (TAD) boundary or contain a CTCF site. Negatively correlated clusters are enriched for bidirectional promoters (Fig.1F).

**Figure 2.**
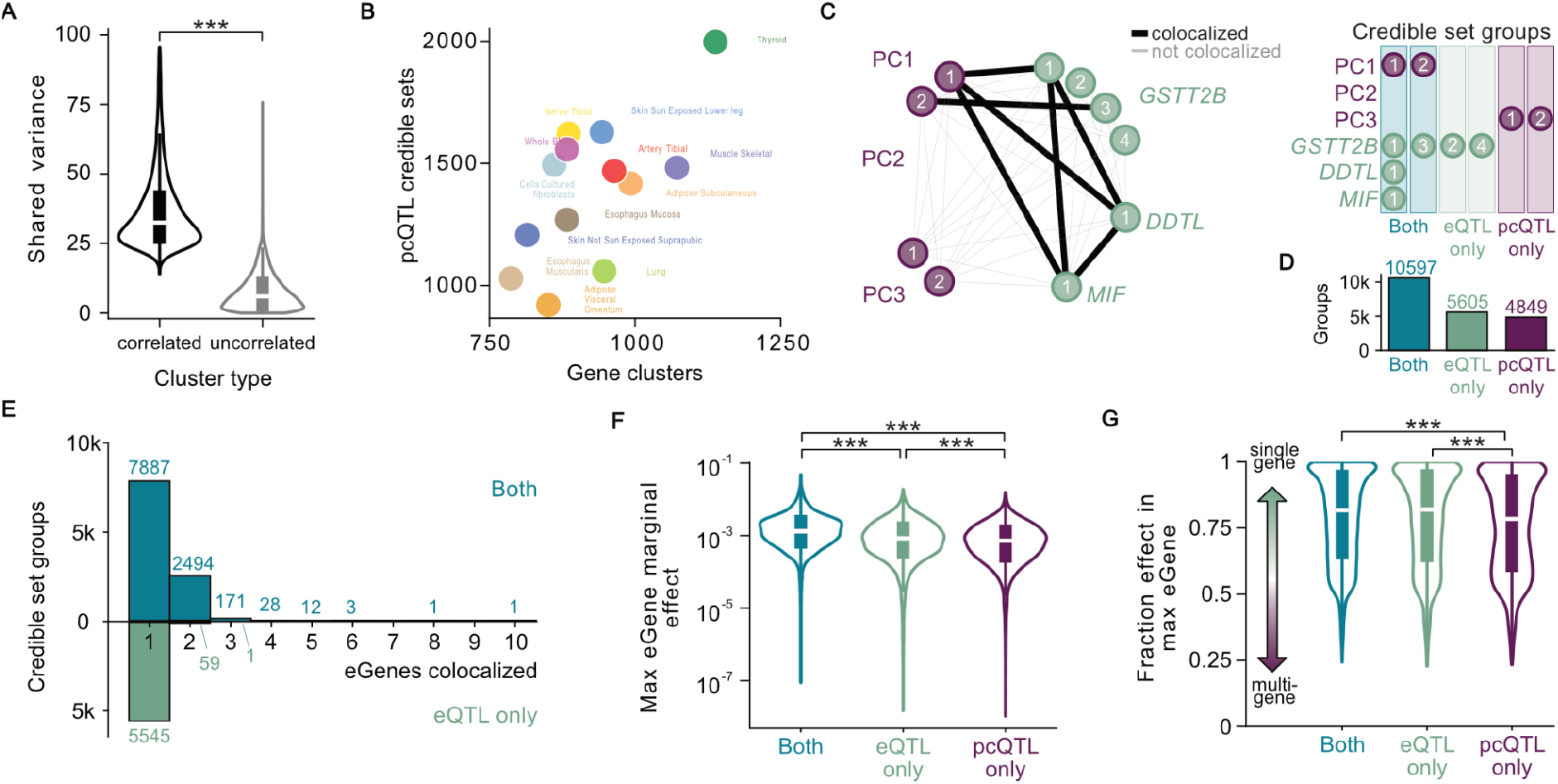
Novel pcQTL discovery. **A** Normalized shared variance (see Methods) explained by PC1 for correlated clusters vs neighboring genes not called as clusters. *** : p<10^−10^ from two-sample t-test. **B** Number of pcQTL credible sets vs number of gene clusters across tissues. **C** Example of credible set groups from a three gene cluster. Nodes are credible sets and edges are colored according to colocalization between the credible sets. **D** Number of credible set groups of each type across all clusters. **E** Number of eGenes colocalized in each credible set group split by whether the group is eQTL only or both a pcQTL and eQTL. **F** The maximum PIP-weighted marginal effect of any credible set in the group on any eGene in the cluster. *** : p<10^−10^ from two-sample t-test. **G** The fraction of effect concentrated into the largest effect eGene: the maximum PIP-weighted marginal effect on any eGene divided by the sum of PIP-weighted marginal effects for all eGenes in the cluster. *** : p<10^−10^ from two-sample t-test.

### A multi-gene QTL framework leverages common variation to detect shared genetic effects

To allow us to jointly consider gene clusters in QTL analysis, we first calculated cluster principal components (PCs) from the normalized gene expression data for each gene cluster (see Methods). As expected for principal component analysis on pre-selected correlated variables, variables have some proportion of shared variance. To summarize the degree to which gene clusters’ expression variance is shared, we calculated the average variance explained across genes in each cluster by PC1, normalized by cluster size such that a cluster with completely correlated gene expression for all genes would have shared variance 100% and a cluster with completely uncorrelated expression would have shared variance 0%. Clusters have significantly higher (p<10^−10^) shared variance (36.6%) than non-correlated neighbor gene pairs (7.6%) (Fig.2A; Supplementary Fig.5). We then use PCs as the dependent variable in QTL mapping (pcQTL). This pcQTL framework allows us to estimate the effect of genetic variants on a shared axis of expression variance across the gene cluster.

We performed pcQTL analysis for all PCs calculated from each gene cluster, and for all variants within a 1Mb-window from any gene in the cluster. Note that in our pipeline, we test all PCs available for a given cluster (e.g. PC 1-5). We then fine-mapped pcQTLs to identify the credible sets of loci driving independent signals, and found between 920-1997 pcQTL credible sets per tissue (Fig.2B; Supplementary Fig.6). For comparison, we also performed single-gene eQTL mapping and fine-mapping for each gene belonging to a cluster, and for the same variant window. In order to identify how many of the pcQTLs are novel, we colocalized each pcQTL credible set to eQTL credible sets for each cluster (Fig.2C). We find that 98% of credible set groups with two or more eGenes are also tagged by a pcQTL (Fig.2E). pcQTL only credible sets that do not colocalize to any eQTL credible sets for any gene in the cluster are considered novel. Across tissues, we find 4859 novel pcQTLs (26.7 % of all pcQTL credible sets) not discovered by any single-gene eQTL analysis (Fig.2D, Supplementary Fig.6).

We then sought to investigate why the pcQTL framework uncovers novel signals. For each novel pcQTL, we calculated the marginal effect of its credible set on the expression of each gene within its cluster (Methods). We found that novel pcQTL signals had both lower maximum marginal effect on any one eGene and lower fraction of total effect concentrated into one eGene (Fig.2F,G, Supplementary Fig.7). This indicates that the pcQTL framework had more power to detect smaller, distributed effects than the single-gene eQTL framework.

### Multi-gene pcQTLs colocalize with new GWAS hits

Most trait-associated variants in genome-wide association studies (GWAS) lie in non-coding regions and are thought to affect traits by regulating gene expression, yet identified eQTLs only colocalize with a small percentage of GWAS hits (43% in GTEx across all tissues) ^11^. We sought to investigate whether our newly-discovered pcQTLs can help explain additional GWAS hits. To do this, we colocalized our pcQTLs with GWAS hits for 74 traits, including cardiometabolic, hematologic, neuropsychiatric, and anthropometric features from UKBB and GIANT ^12^. For comparison, we also colocalized GWAS hits and eQTLs for each gene in each cluster. Using single-gene eQTL mapping alone, 1570 GWAS hits could be linked to at least one single-gene eQTL in a cluster in a given tissue (representing 633 unique GWAS hits across tissues). With multi-gene mapping on cluster PCs, an additional 535 GWAS hits were colocalized with a pcQTL but not with any single-gene eQTLs with a cluster in a given tissue (representing 142 unique GWAS hits) (Fig.3A). This represents a 34% increase in colocalizations (22% increase in unique GWAS hits) compared with single-gene methods alone.

**Figure 3.**
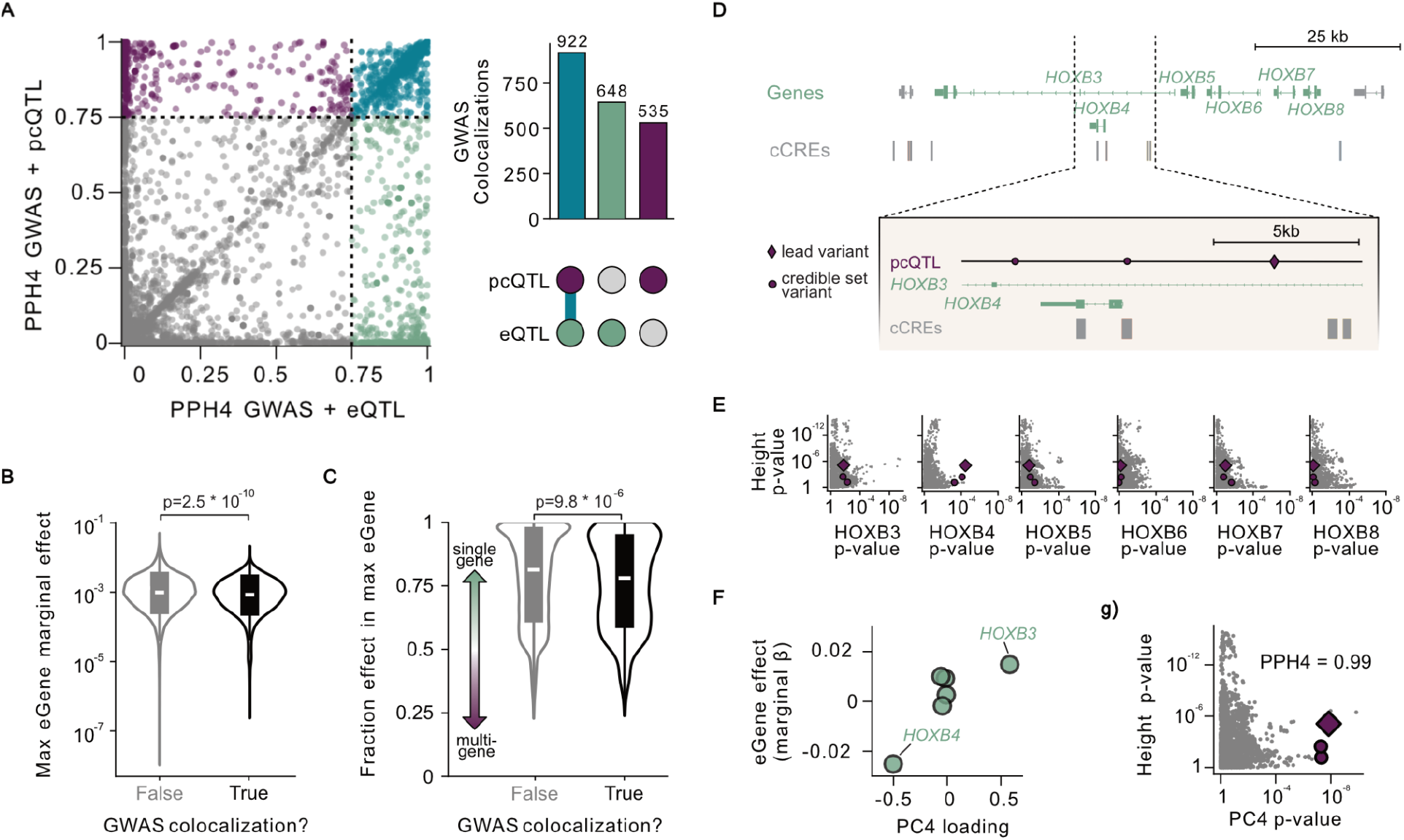
pcQTL colocalization with complex trait associated variants. **A** Maximum posterior probability of a shared causal variant (PPH4) underlying the GWAS hit and any pcQTL credible set for the cluster vs for any eQTL credible set in the cluster. PPH4=0.75 is shown as a dotted line. Upset plot: total GWAS colocalizations across all clusters and tissues. **B** The maximum PIP-weighted marginal effect on any eGene in the cluster, split by whether or not the credible set group colocalizes with a GWAS hit. P-value from two-sample t-test. **C** The fraction of effect concentrated into the largest effect eGene: the maximum PIP-weighted marginal effect on any eGene divided by the sum of PIP-weighted marginal effects for all eGenes in the cluster, split by whether or not the credible set group colocalizes with a GWAS hit. P-value from two-sample t-test. **D** *HOXB3, HOXB4, HOXB5, HOXB6, HOXB7, HOXB8* gene cluster and cCREs in esophagus muscularis, Close-up of region with fine-mapped novel PC4 pcQTL credible set. **E** Nominal p-values for GWAS hit and each eGene in the cluster independently, with the PC4 pcQTL credible set variants highlighted. **F** Marginal effect of pcQTL on each eGene vs PC4 loading onto each eGene. **G** Nominal p-values for height GWAS and PC4. Posterior probability of colocalization between the GWAS hit and the pcQTL credible set PPH4 is 0.99.

Consistent with evidence that large-effect QTL variants are less likely to colocalize with GWAS hits because stronger regulatory effects at crucial genes tend to be purged by negative selection ^13^ we found that larger-effect QTLs in our analysis were less likely to colocalize with GWAS hits (t-test p=2.5e-10) (Fig.3B). In contrast, variants whose effects were distributed across multiple genes (lower fraction maximum effect) showed higher rates of colocalization (t-test p=9.8e-6), precisely the type of signal that pcQTL mapping can capture more effectively (Fig.2G, Fig.3C). The association between fraction of effect and likelihood to be colocalized with a GWAS hit remained significant (p=2.1e-5) even in a regression analysis with maximum effect size as a covariate.

An example of one such novel pcQTL credible set colocalization illustrates how PCs can boost power by summarizing distributed effects across genes. The colocalization is between PC4 for a 6-gene *HOXB* cluster (*HOXB3, HOXB4, HOXB5, HOXB6, HOXB7*, and *HOXB8)* in esophagus muscularis tissue and a GWAS hit for height (Fig.3D). Although the *HOXB* transcription factors play a well established role in development ^14^ and single-gene eQTL analysis maps a eQTL for *HOXB3*, this eQTL did not colocalize with the height GWAS hit (posterior probability of colocalization-PPH4=4.7e-5). When we instead used PC4 for pcQTL mapping, we detected an additional locus which colocalized with the GWAS hit (PPH4=0.99) (Fig.3G). Fine-mapping established a three variant credible set for the pcQTL (Fig.3D). The credible set variants have below-threshold nominal p-values for some genes individually, but none reach significance on their own (Fig.3E).

One variant of the credible set overlaps a candidate *cis*-regulatory element (cCRE) region for esophagus muscularis ^15^ (Fig.3D). Based on PC loadings, PC4 primarily captures inverse variation between *HOXB3* and *HOXB4* (Fig.3F). Notably, enhancer-promoter links from the ABC model indicates the cCRE acts as both the promoter of *HOXB4* and an enhancer for *HOXB3* ^16^. The two genes are positively correlated (Fig.1E) but this positive correlation is summarized in the first 3 PCs, allowing PC4 to capture a subtler inverse effect in the cCRE and map a novel pcQTL which colocalizes with the GWAS hit.

Another novel pcQTL credible set colocalization demonstrates a case where the combined impact of multiple genes, rather than either gene independently, may be responsible for an observed complex trait association. A pcQTL from PC2 for a two-gene cluster in subcutaneous adipose tissue containing NLRC3 and CLUAP1 colocalized with a GWAS hit for BMI (PPH4=0.930) and body fat percentage (PPH4=0.751) (Supplementary Fig.8). Each of these genes could plausibly impact BMI individually. Higher NLRP3 plasma expression has been correlated with higher body weight ^17,18^. RNAi targeting of CLUAP1 in Drosophila showed significant body weight increase, but KO in mice did not ^19^. However, CLUAP1 is part of the intraflagellar transport complex B (IFT-B) required for cilia biogenesis, and other cilia related proteins have been implicated in adipocyte differentiation and obesity giving CLUAP1 a plausible mechanism of action on BMI ^20^. While single-gene eQTL analysis does pick up eQTLs regulating NLRC3 and CLUAP1 individually, none of these single-gene effect loci colocalize with the BMI or body fat GWAS hits (max PPH4=1.01e-3). However, there is a significant difference (Fisher-Z p-value=4.4e-4) in the Spearman correlation between the two genes depending on genotype at the lead variant of the pcQTL loci (ρ=-0.06 for the homozygous reference vs ρ=-0.36 for the homozygous alternate) indicating the potential importance of a combined effect between the two genes captured by PC2.

## Discussion

eQTL studies have focused on single gene at a time analyses despite evidence of molecular pleiotropy, where variants impact the molecular function of multiple genes. To detect such pleiotropic variants, previous work has primarily focused on mapping single-gene eQTLs and reporting instances where a variant is independently associated with multiple genes rather than leveraging shared effects ^1^. We demonstrate that when we move from considering genes as discrete functional units to jointly considering multiple neighboring genes, we detect novel QTLs and new colocalizations with complex-trait-associated variation.

In our study, pcQTLs represent a proof-of-concept methodology for the joint consideration of neighboring genes in eQTL mapping. Other ways to cluster genes, additional dependent variables constructed from combinations of PCs, as utilized for *trans*-QTL mapping in ^21^, or conceptually similar methods using multi-trait fine-mapping ^22,23^ could also be adapted to utilize shared local gene regulation to discover additional signals. Further work will be required to determine which clusters of genes and which statistical tools are best suited to different aspects of QTL discovery. However, the dramatic increase in discoveries and complex trait-associated variation even with a methodologically straightforward approach as pcQTLs highlights that leveraging shared *cis*-regulation and subsequent co-expression of neighboring genes is an important aspect of QTL mapping that can elucidate new trait and disease biology. Combined, this approach recognizes the complexity of gene regulation, leveraging the expression correlations among nearby genes to capture novel signals of molecular pleiotropy. However, as molecular pleiotropy can describe both local and distal effects, we classify variants with effects on multiple, proximal genes as having molecular “proxitropy”. We expect future studies will benefit by moving from a classical genetics view of a causal variant impacting a single causal gene to one integrating the complexity of local regulation.

## Methods

### Processing of GTEx RNA-seq data

Normalized expression data from GTEx v8 ^24^ for adipose (subcutaneous), adipose (visceral omentum), tibial artery, cultured fibroblasts, esophagus (mucosa), esophagus (muscularis), lung, skeletal muscle, tibial nerve, skin (not sun exposed), skin (sun exposed), thyroid, and whole blood were downloaded from the GTEx portal. Tissues were chosen based on having the largest sample sizes (N>400). Expression data was then residualized on 60 PEER factors, the top 5 genotype PCs, sequencing platform, sequencing protocol, and sex. These were the same covariates that were used in the standard GTEx eQTL pipeline ^11^.

### Calling clusters from RNA-seq data

To call clusters, a stepwise, iterative “sliding window” approach was used to identify stretches of neighboring, co-expressed genes (“gene clusters”). First, gene expression Spearman correlation matrices were calculated for each chromosome separately, using the normalized and residualized gene expression. Correlations significant at p < 0.05 after Bonferroni correction (accounting for the total number of genes on the chromosome) were considered significant. Positive and negative correlations were both considered. We identified clusters where more than 70% of pairwise-correlations between genes within the cluster were significant. To find these clusters, we started with a maximum window size of n=50 genes and continued until a minimum window size of n=2 genes. Algorithmically, assuming N genes total on the chromosome, we:

1. Check if the first 0-n neighboring genes on the chromosome are sufficiently correlated and do not belong to any existing clusters. If so, we record those n genes as a cluster.
2. Check if the next 1-n+1 neighboring genes are sufficiently correlated and do not belong to any existing clusters. If so, we record these as a cluster.
3. Continue with 2-n+2, 3-n+3, …, N-n, N. As we move the window over one gene at each step, we check if the genes are sufficiently correlated and do not belong to any existing clusters. If so, we record them as a cluster.
4. Reduce the cluster size n to n-1 and restart at step 1. We repeat this process until the minimum window size (n=2) is reached.

This approach captures genes in the largest possible cluster with significant within-cluster correlations. As false positives correlations due to cross-mappable reads were a concern, we compared the number of QTLs mapped for clusters with and without any genes containing cross-mappable 75-mers (Supplementary Fig.4) ^25^. As the QTL discovery rate was the same for cross-mappable and non-cross mappable genes, we continued to consider cross-mappable clusters for further analysis.

### Computing cluster enrichments for various annotations

In order to evaluate whether clusters were enriched for particular annotations (such as “paralogs” or “shared GO terms”), we calculated cluster enrichments for all clusters against a background of null clusters. To do this, we assigned correlated and null clusters a binary label for whether or not they belonged to several (not mutually exclusive) categories. These labels were used as the dependent variable in logistic regression to calculate enrichment odds ratios, with the number of genes in the cluster as a covariate. Null clusters were of all sets 2, 3, 4, or 5 of neighboring genes in each tissue that were not part of a correlated cluster in that tissue. The number of genes in the cluster was then included as a covariate in all regressions. Because many annotations (i.e. having a CTCF site between the genes’ starts) would be influenced by the size of the cluster, we also wished to account for cluster size in our enrichment analysis. Cluster size followed a bimodal distribution with gene-start to gene-start peaks at < 1kb (shared promoters) and 50kb (Supplementary Fig.4). Because of its non-normality, cluster size could not be included as a covariant for regressions. Instead, for each cluster size separately, we re-sampled the null distribution for that cluster size to match the minimum gene-start to gene-start distance (between any two cluster genes) distribution for correlated clusters at that cluster size. This resampled null was used for most regressions. The non-resampled null was used for annotations directly concerning the distance between TSSs (bidirectional promoter or shared promoter).

For each correlated and null cluster, we annotated whether each belonged to the following categories:

- <u>Paralogs</u>: if any two (or more) genes in the cluster were listed as paralogs. Parlog information was obtained from the biomart webtool for ensemble 97 ^26^.
- <u>Shared GO term</u>: if any biological process (BP) GO term was shared between any two (or more) genes. GO terms for each gene were obtained from the biomart webtool for ensemble 97 ^26^.
- <u>Bidirectional promoter</u>: if the 5’ end of any annotated GENCODE v26 ^27^ transcript for gene A was within 1000 bp of the 5’ end of any annotated transcript for gene B, for any pair of genes A/B in the cluster.
- <u>Enhancer</u>: If in ABC enhancer-gene predictions for a manually curated cell-type matched to the GTEx tissue (Supplementary Data 1), a pair of genes are both listed as sharing an ABC enhancer ^16^.
- <u>Cross CTCF peak</u>: CTCF ChiP-seq run on GTEx samples from ENTEx ^28^ was used. For each tissue, CTCF peaks from a matching experiment were downloaded from ENCODE (Supplementary Data 2). If a CTCF peak fell between the window defined by the outer edges of any transcript in the cluster, that cluster was annotated as crossing a CTCF peak.
- <u>Cross transcriptionally associated domain (TAD) boundary</u>: TAD boundaries calculated from Hi-C data in GM12878 with directionality index (DI) at a 10kb resolution were downloaded from TADKB ^29^ and converted to hg38 with liftOver^30^ If the edge of a TAD fell between the window defined by the outer edges of any transcript in the cluster, that cluster was annotated as crossing a TAD boundary.

### Calculating cluster principal components (PCs) and normalized shared variance variance

For each cluster of genes, PCA (sklearn v1.3.2) was used to find a shared axis of expression variance. Principal components were constructed as a linear combination of the normalized, residualized expression of the genes in the cluster.

To find the shared variance explained by PC1, given an cluster of *n* genes, we first summed the variance explained for each gene in the cluster by PC1:

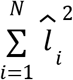

where the variance explained by PC1 for gene *i* is the squared loading 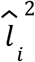 for PC1 onto that gene. We then subtract 1 and normalized to cluster size, so that for a cluster of *N* genes, shared variance is

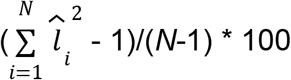

This rescales the value so that regardless of cluster size, if all variables are uncorrelated, the normalized shared variance is 0 and if all variables are completely correlated the normalized shared variance is 100.

### Mapping pcQTLs and eQTLs

QTLs were mapped on all variants within 1Mb of the minimum starting boundary and the maximum ending boundary of all genes in a given cluster for both pcQTLs and eQTLs with tensorQTL ^31^. This ensured that both eQTL and pcQTL mapping was applied to the same set of variants. Input phenotypes were:

- eQTL mapping: the expression of each gene in each cluster
- pcQTL mapping: the PCs on the expression of the gene cluster

TensorQTL was run with default settings in *cis* mode to get phenotype-level summary statistics with permutation based empirical p-values for each phenotype and *cis_nominal* mode to get summary statistics for all variant-phenotype pairs. To finemap, SuSiE ^32^ was used with default settings to finemap independent loci to 95% credible sets of variants for all expression and PC phenotypes.

### PIP-weighted marginal effects

To investigate the extent to which pcQTL influenced each gene within a target cluster individually, we calculated PIP-weighted marginal effects. The marginal effect of a given pcQTL credible set on the expression of each gene within its cluster was calculated by taking a PIP-weighted average across all variables in the credible set. For each credible set variant and for each cluster gene, we multiplied the nominal effect size β of that variant on the cluster genes by the PIP probability. These PIP-weighted effects were then summed and divided by the sum of the PIP probabilities. This allowed us to compare the effects of pcQTL credible sets on gene expression in expression coordinate space.

The extent to which a given credible set had its efßfect concentrated in just one gene or spread across multiple genes was quantified with the fraction of total effect quantified by the single largest effect gene. For a credible set on a cluster of *n* genes, impacting each gene *i* with squared marginal effect 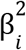, given that 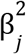 is the largest, we calculated

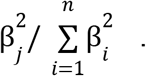

### PIP-weighted variant annotations

Ensembl Variant Effect Predictor (Ensembl VEP v114) ^33^ was used to annotate each variant with a likely function. Variant annotations were converted to probabilities for credible sets. To evaluate whether a credible set was enriched for a particular annotation, for each annotation, the PIP values for all variants in the credible set with the annotation were summed and then divided by the total of all the PIPs. These probabilities were used as the dependent variable in logistic regression to calculate odds ratios.

### Colocalization of pcQTL, eQTL and GWAS hits

We used coloc-SuSiE (v5) ^34^ to colocalize pcQTL, eQTL, and GWAS signals. GWAS summary stats from a publicly available resource of 114 GWAS (for 74 distinct traits, including cardiometabolic, hematologic, neuropsychiatric, and anthropometric features from UKBB and GIANT) harmonized and imputed to GTEx variants were used ^12^. For each cluster, all eQTL, pcQTL, and GWAS credible sets were colocalized with all other eQTL, pcQTL, and GWAS credible sets. An undirected graph was constructed with each credible set as a node and an edge connecting two nodes if the probability of colocalization (ppH4) between those credible sets was > 0.75. Credible set groups were the set of connected components of the graph.

## Data availability

All processed GTEx data are available via GTEx portal (https://www.gtexportal.org/home/downloads/adult-gtex). Paralog and GO terms are available on biomart (https://www.ensembl.org/info/data/biomart/index.html). CTCF peaks are available on the ENCODE portal (https://www.encodeproject.org/). TAD boundaries are available on TADKB (http://dna.cs.miami.edu/TADKB/). ABC predictions across cell types are available from the Engreitz lab (https://www.engreitzlab.org/resources). cCREs are available from SCREEN (https://screen.encodeproject.org/). GWAS summary stats are available on Zenodo (https://zenodo.org/records/3629742#.Y9rTQOzMIUF). Clusters and summary stats for pcQTLs are available on Zenodo (https://doi.org/10.5281/zenodo.15605351).

## Code availability

TensorQTL was used for eQTL and pcQTL mapping and is available at https://github.com/broadinstitute/tensorqtl. coloc was used for colocalizations (https://chr1swallace.github.io/coloc/index.html). Scripts used to call clusters, pipeline for data processing, and notebooks to generate all figures are available at https://github.com/kal26/pc_qtls.

## Acknowledgements

We thank the donors and their families for their generous gifts of biospecimens to the GTEx research project. The Genotype-Tissue Expression (GTEx) project was supported by the Common Fund of the Office of the Director of the National Institutes of Health (http://commonfund.nih.gov/GTEx). Additional funds were provided by the National Cancer Institute (NCI), National Human Genome Research Institute (NHGRI), National Heart, Lung, and Blood Institute (NHLBI), National Institute on Drug Abuse (NIDA), National Institute of Mental Health (NIMH), and National Institute of Neurological Disorders and Stroke (NINDS). This research was supported by National Institutes of Health grants R01MH12524, U01AG072573, U01HG012069 to S.B.M. K.L. is supported by the Stanford Genome Training Program (SGTP; NIH/NHGRI T32HG000044). T.G. is supported by the Knight-Hennessy Scholars fellowship. The funders had no role in study design, data collection and analysis, decision to publish or preparation of the manuscript.

## Author contributions

S.B.M., T.G., and K.L conceived and designed the study. T.G. designed the cluster calling algorithm. K.L, performed most of the analysis. S.B.M., T.G., and K.L wrote the manuscript.

## Extended Data

**Extended Data Figure 1.**
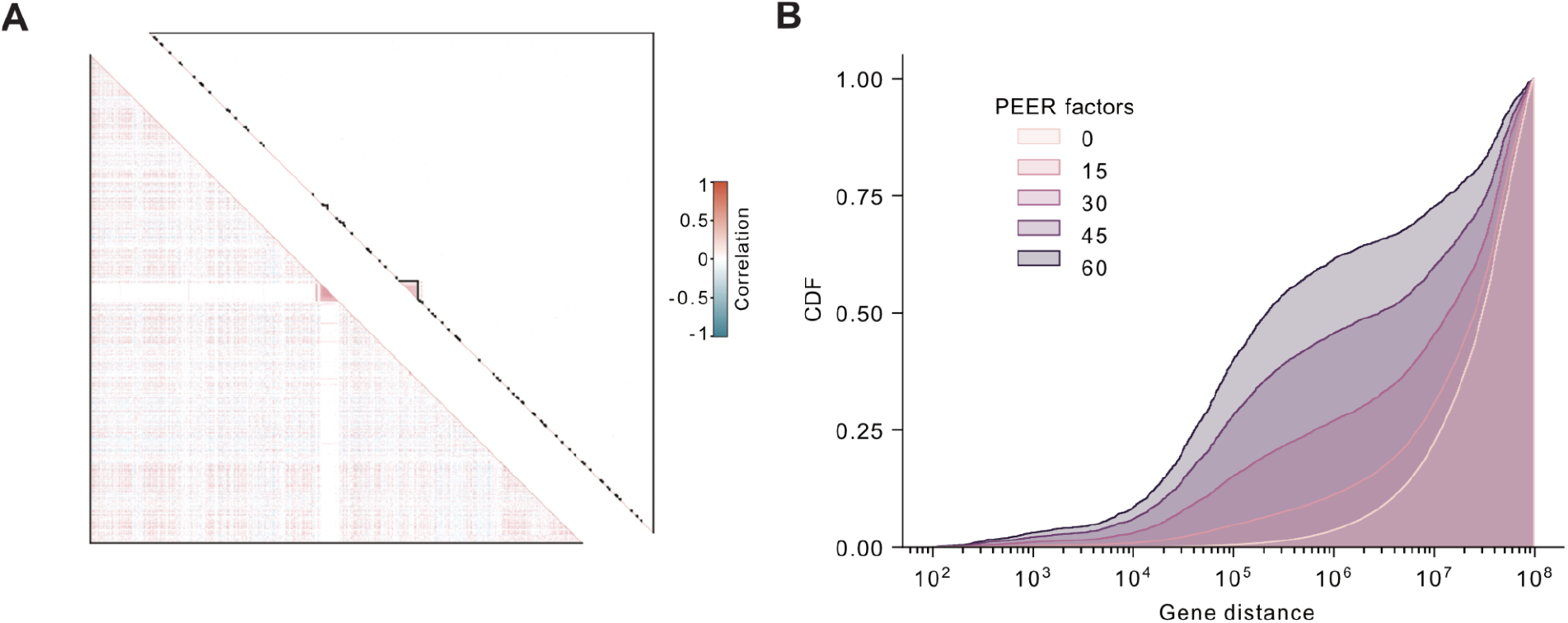
PEER factor residualization’s effect on correlation. **A** Correlation matrix of gene expression in chromosome 17 for sun exposed skin; Spearman correlation before (lower) and after (upper) residualization with 60 PEER factors. Correlations with Bernoulli significant p-value are shown. **B** CDF of pairwise gene distances of genes with significantly correlated gene expression profiles after residualization with 0, 15, 30, 45, and 60 PEER factors.

**Extended Data Figure 2.**
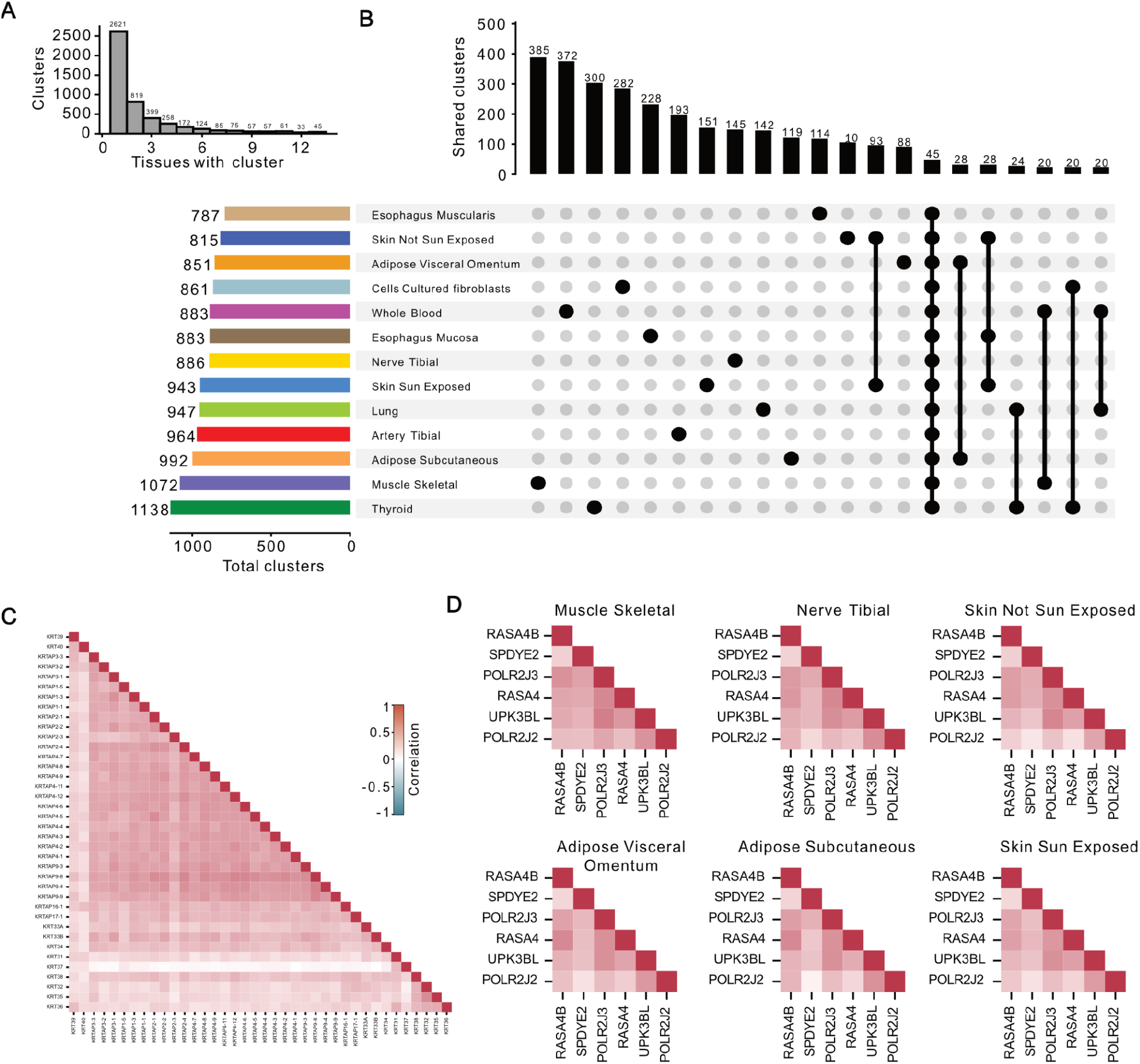
Tissue specificity of clusters. **A** Histogram of the number of tissues a cluster is in. **B** Upset plot detail of which tissues a cluster is shared across for all tissue combinations with 20 or more clusters. **C** A tissue-specific thirty-eight gene cluster on chromosome 17 from sun exposed skin gene expression. Color is the expression Spearman correlation. **D** A six gene cluster on chromosome 6, shared across 6 tissues. Color is the expression Spearman correlation.

**Extended Data Figure 3.**
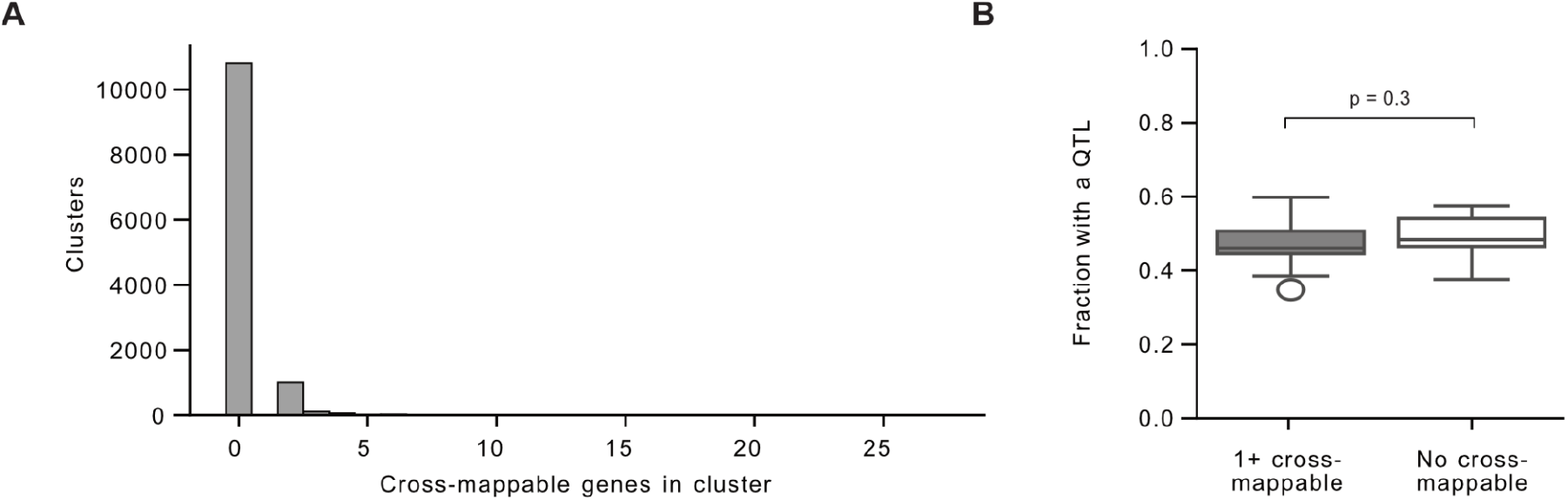
Cross-mappability of clusters. **A** Distribution of minimum gene-start to gene-start distance for all clusters. **B** Gene clusters were called on expression data for cultured fibroblasts for each chromosome on expression data with gene locations shuffled.

**Extended Data Figure 4.**
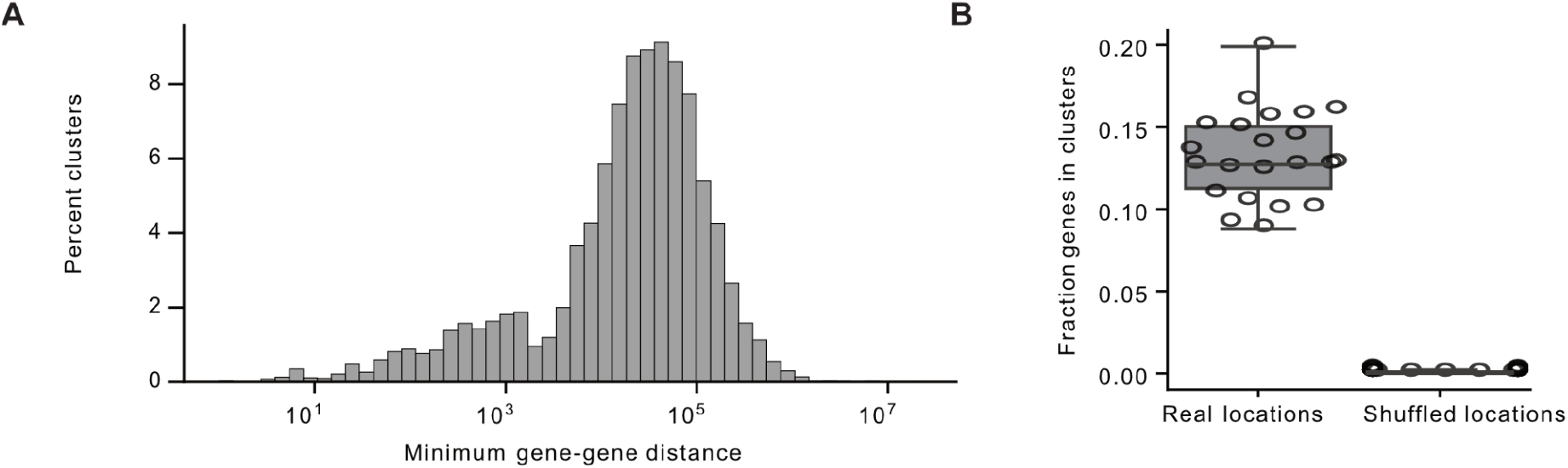
Cluster size distribution and shuffled null. **A** Histogram of the number of genes in each cluster with more than 100 cross-mappable 75-mers. **B** Fraction of gene clusters with at least one significant QTL for clusters with at least one cross-mappable gene pair or with no cross mappable gene pairs. P-value from independent t test.

**Extended Data Figure 5.**
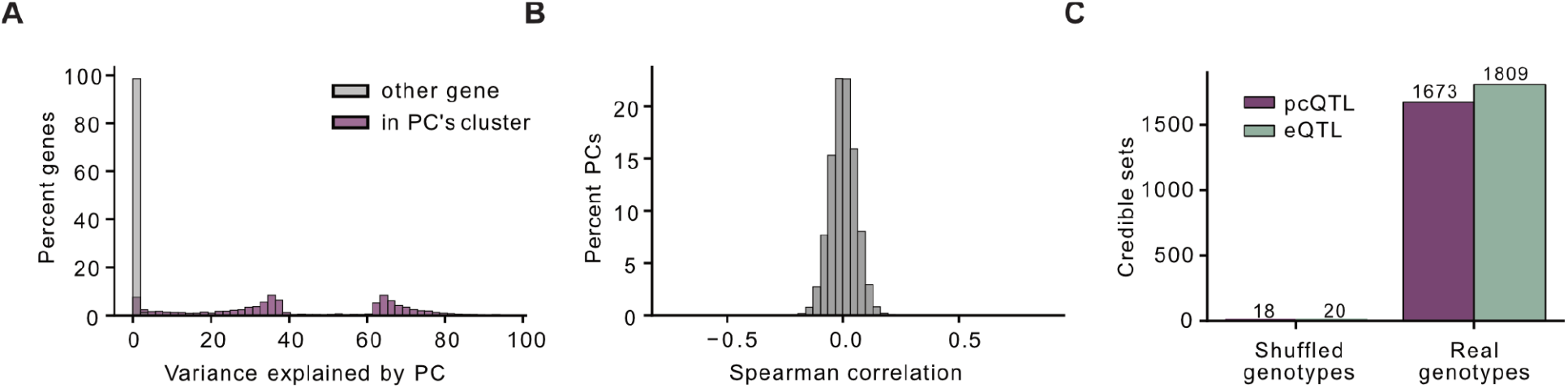
Local PC properties. **A** Variance explained by each PC phenotype for all genes split by whether or not the gene is in the cluster whose expression was used to calculate the PC. **B** Spearman correlation each PC to all other PCs for all cultured fibroblast PCs. **C** pcQTLs and eQTLs were called for a genotype-shuffled null. The number of significant credible sets for pcQTLs and eQTLs for cultured fibroblasts for the original genotypes and the shuffled null is shown.

**Extended Data Figure 6.**
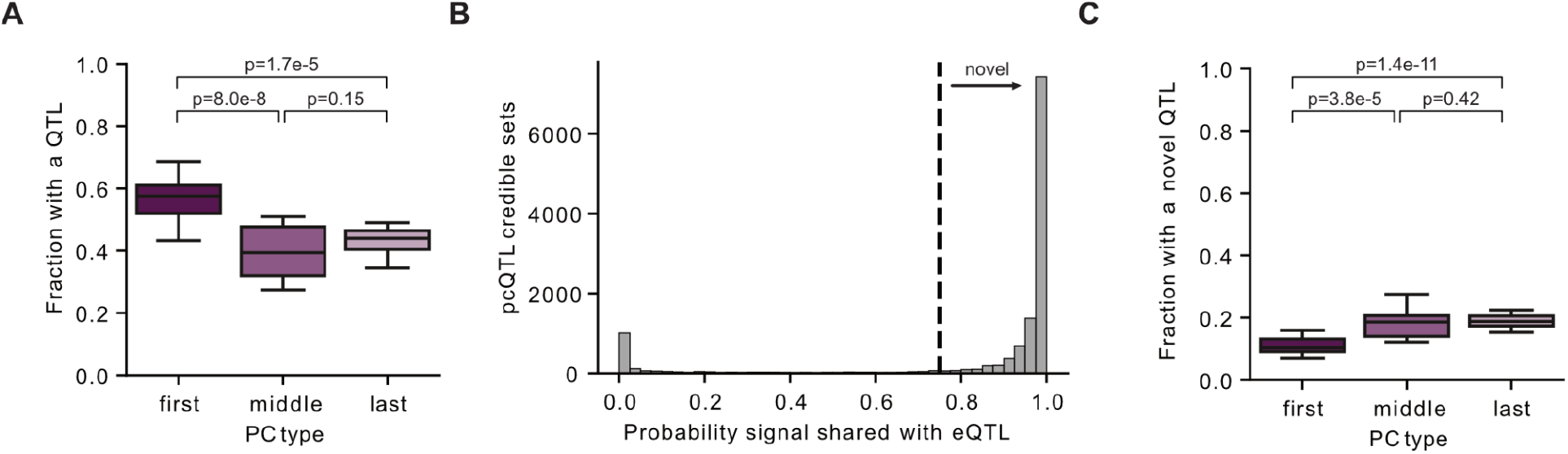
pcQTL discoveries by PC order. **A** The fraction of PC phenotypes across tissues with at least one significant QTL split into the primary (first) PC, the last PC, and all other PCs (middle). p-values are from a paired-sample t-test. **B** Distribution of the maximum PPH4 colocalization probability for each pcQTL credible set with any cluster eQTL credible set. Dotted line PPH4=0.75. **C** The fraction of PC phenotypes across tissues with at least one novel significant QTL split into the primary (first) PC, the last PC, and all other PCs (middle). p-values are from a paired-sample t-test.

**Extended Data Figure 7.**
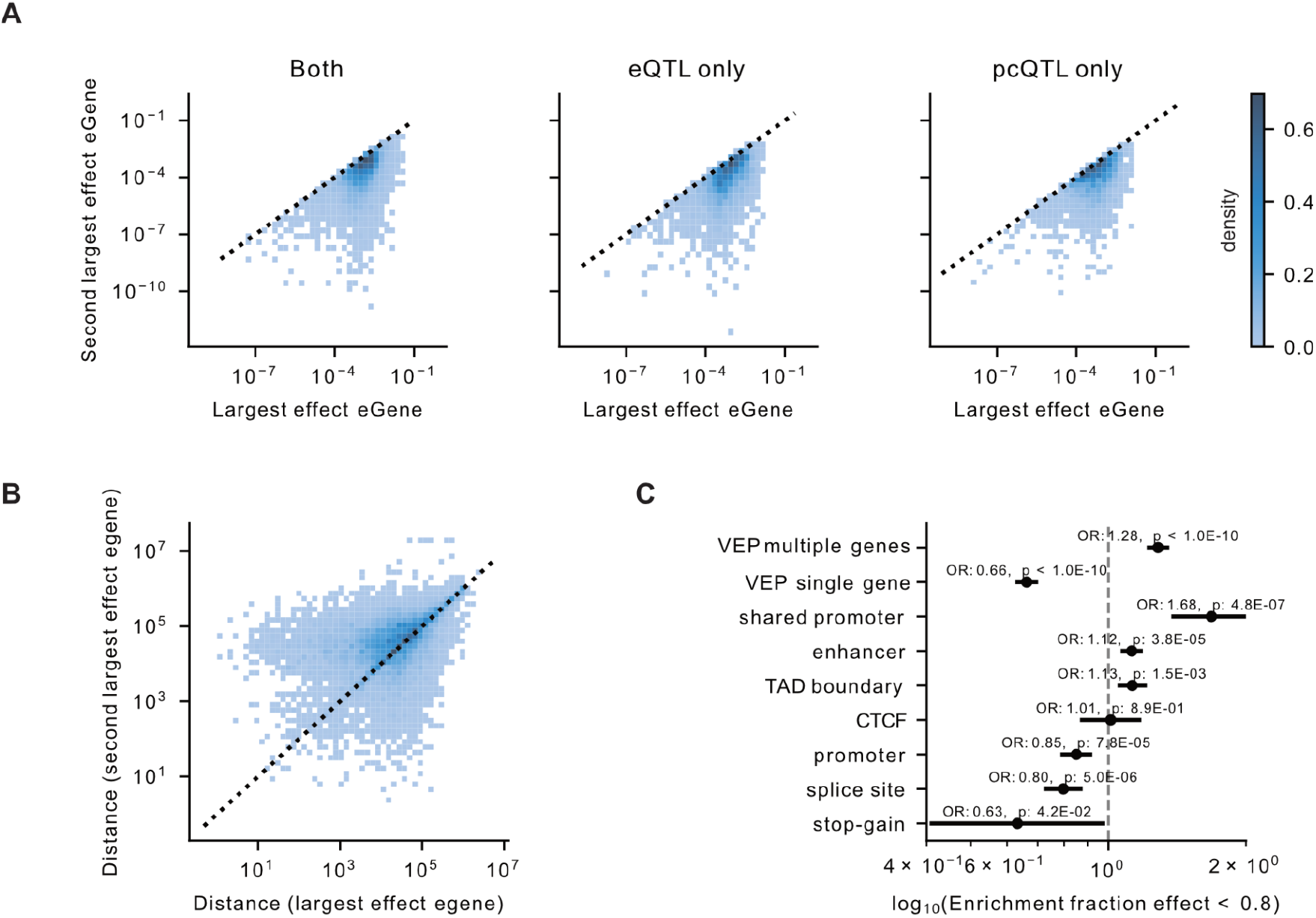
Distribution of eGene marginal effects. **A** The second largest PIP-weighted effect of a QTL on any eGene in a cluster vs the largest PIP-weighted effect of a QTL on any eGene in a cluster, split by if the QTL is discovered as both a pcQTL and eQTL, an eQTL only, or a pcQTL only. **B** The distance from the lead variant of the QTL credible set to the gene start of the second largest effect eGene vs the largest effect eGene. **C** Annotation enrichments for PIP-weighted variant effect predictor categories for QTL credible sets split by if the fraction of total effect quantified by the single largest effect gene is higher or lower than 0.8.

**Extended Data Figure 8.**
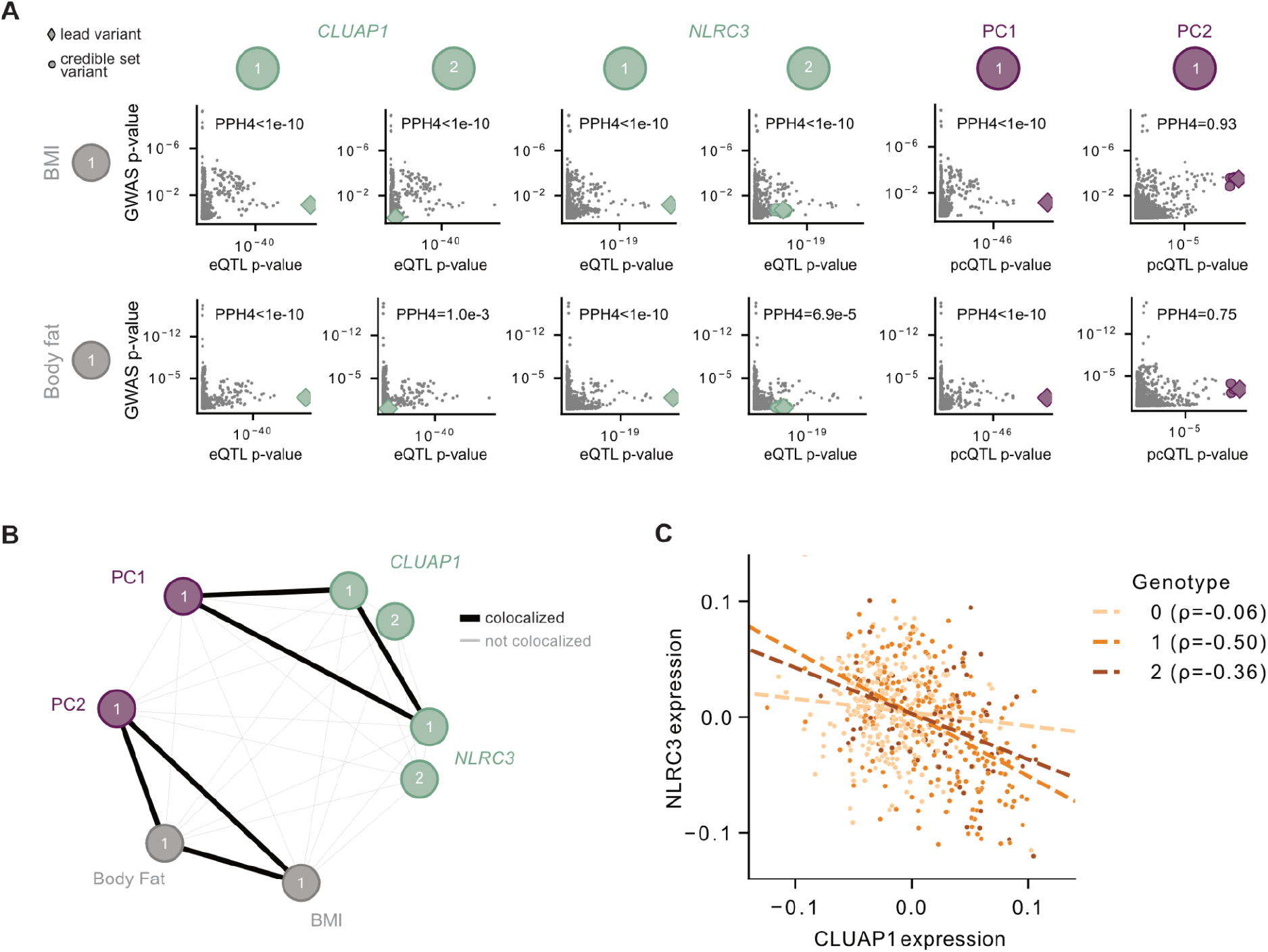
BMI colocalization for a NLRC3 and CLUAP1 pcQTL. **A** GWAS variant nominal p-values for BMI and body fat vs QTL nominal p-values for each gene and PC. Columns are credible sets, highlighted variants are the fine-mapped credible set variants. which correspond to highlighted variants in plots. PPH4 is probability of colocalization between the GWAS hit and the pcQTL credible set. **B** Credible set group graph for the cluster. **C** normalized expression for NLRC3 vs normalized expression for CLUAP1 with points colored by the individual’s genotype. Lines are best fit from linear regression and ρ is Spearman ρ.

## Supplementary Information

**Supplementary Table 1.**
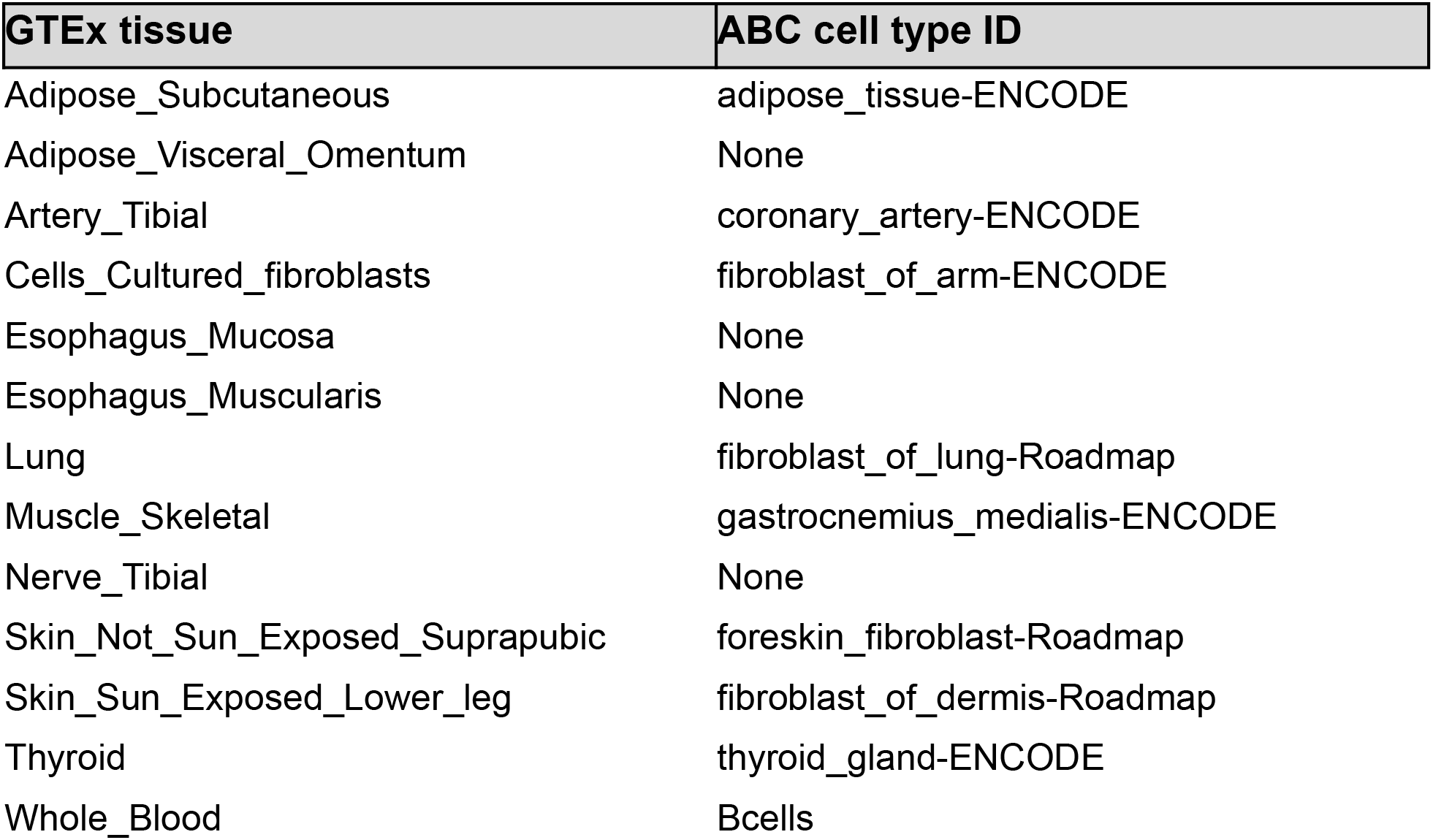
GTEx tissue matches for ABC enhancer-gene cell-types.

**Supplementary Table 2.** GTEx tissue matches for ENCODE files for CTCF peaks.

| **GTEX tissue** | **ENCODE file ID** |
| --- | --- |
| Adipose_Subcutaneous | ENCFF173APP |
| Adipose_Visceral_Omentum | ENCFF051ZVS |
| Artery_Tibial | ENCFF508XDM |
| Cells_Cultured_fibroblasts | ENCFF130ISX |
| Esophagus_Mucosa | ENCFF862AXF |
| Esophagus_Muscularis | ENCFF862AXF |
| Lung | ENCFF786YIA |
| Muscle_Skeletal | ENCFF721RAM |
| Nerve_Tibial | ENCFF831KJJ |
| Skin_Not_Sun_Exposed_Suprapubic | ENCFF525JGP |
| Skin_Sun_Exposed_Lower_leg | ENCFF405VLP |
| Thyroid | ENCFF498GQK |
| Whole_Blood | ENCFF796WRU |

